# Polycomb Repressive Complex 2 promotes atherosclerotic plaque vulnerability

**DOI:** 10.1101/2024.12.02.626505

**Authors:** Divyesh Joshi, Raja Chakraborty, Tejas Bhogale, Jessica Furtado, Hanqiang Deng, James G. Traylor, Anthony Wayne Orr, Kathleen A. Martin, Martin A. Schwartz

## Abstract

**Key findings:**

1. PRC2 regulates EC shear stress responses.
2. PRC2 governs Klf2/4 suppression downstream of Pcdhg.
3. High PRC2 in ASCVD-prone arterial regions suppresses Klf2/4 to promote ASCVD.
4. Athero-protective Klf2/4 induction upon PRC2 inhibition requires Notch signaling.
5. Tazemetostat, an FDA approved PRC2 inhibitor, slows ASCVD progression and improves markers of plaque stability.

Atherosclerotic cardiovascular disease (ASCVD), the leading cause of mortality worldwide, is driven by endothelial cell inflammatory activation and counter-balanced by anti-inflammatory transcription factors Klf2 and Klf4 (Klf2/4). Understanding vascular endothelial inflammation to develop effective treatments is thus essential. Here, we identify, Polycomb Repressive Complex (PRC) 2, which blocks gene transcription by trimethylating histone3 Lysine27 in gene promoter/enhancers, as a potent, therapeutically targetable determinant of vascular inflammation and ASCVD progression. Bioinformatics identified PRC2 as a direct suppressor of Klf2/4 transcription. Klf2/4 transcription requires Notch signaling, which reverses PRC2 modification of Klf2/4 promoter/enhancers. PRC2 activity is elevated in human ASCVD endothelium. Treating mice with established ASCVD with tazemetostat, an FDA approved pharmacological inhibitor of PRC2, slowed plaque progression by 50% and drastically improved markers of plaque stability. This study elucidates a fundamental mechanism of vascular inflammation, thus identifying a potential method for treating ASCVD and possibly other vascular inflammatory diseases.

## 1. Introduction

ASCVD, the leading cause of death globally, arises as a consequence of converging metabolic, inflammatory, and biomechanical risk factors^1,2^. Endothelial cells (ECs) lining the vasculature transduce the fluid shear stress (FSS) from blood flow and into biochemical responses that ultimately control gene transcription and thus cell phenotype. In straight regions of arteries, laminar shear stress (LSS) from blood flow protects from ASCVD by upregulating Kruppel like factors 2 and 4 (Klf2/4) in ECs. Conversely, disturbed shear stress (DSS) at sites of vessel branching or curvature (modeled in vitro by oscillatory shear stress or OSS) predisposes these regions to plaque formation by suppressing Klf2/4 and activating the pro-inflammatory NF-κB pathway^1,2^. These two pathways are mutually inhibitory, the balance between these pathways plays a crucial role in determining disease susceptibility and progression^3–7^. Klf2 and Klf4 are co-regulated and share many gene targets in ECs, with substantial functional redundance. Consistent with their protective function, increasing or decreasing EC Klf2/4 expression decreases or increases, respectively, the incidence and severity of vascular inflammation and disease in multiple mouse models^8,9^. However, our limited knowledge of shear stress-dependent Klf2/4 transcriptional regulation has limited the development of clinically effective methods to elevate Klf2/4 for treatment of ASCVD.

Towards this goal, our genome-wide CRISPR-Cas9 screen assessed hemodynamic regulation of Klf2/4 transcription, identifying positive contribution from mitochondrial metabolism and the Notch pathway^9,10^. A study focused on Klf2/4 suppressors identified a pro-atherogenic role of the protocadherin gamma (Pcdhg) gene family, which suppresses Klf2/4 transcription by blocking its induction through the Notch pathway^9^. In this study, we identified PRC2 as a key regulator of endothelial inflammatory gene expression downstream of Pcdhg and explored its efficacy as a target for treating atherosclerosis.

## 2. Results

### Polycomb Repressive Complex 2 (PRC2) in endothelial responses to FSS

We initially profiled EC gene expression under LSS, to model physiological protective blood flow and under OSS, to model ASCVD-associated pathological blood flow. Analysis of differentially expressed genes (DEGs) (Fig 1a) showed the expected elevation of Klf2, Klf4 and their target gene Nos3 under LSS while the inflammatory mediator E-selectin (Sele) was high under OSS (Fig S1a), confirming normal responses to these FSS patterns. Likewise, pathway/process analysis showed overrepresentation of shear stress and atherosclerosis categories (Fig 1b). Upstream regulator analysis of DEGs using EnrichR^11^ identified the core PRC2 component SUZ12 as the leading mediator (Fig 1c), with the PRC2 catalytic subunit EZH2 as an additional strong hit. PRC2 may thus be a key node governing EC response to FSS.

**Figure 1.**
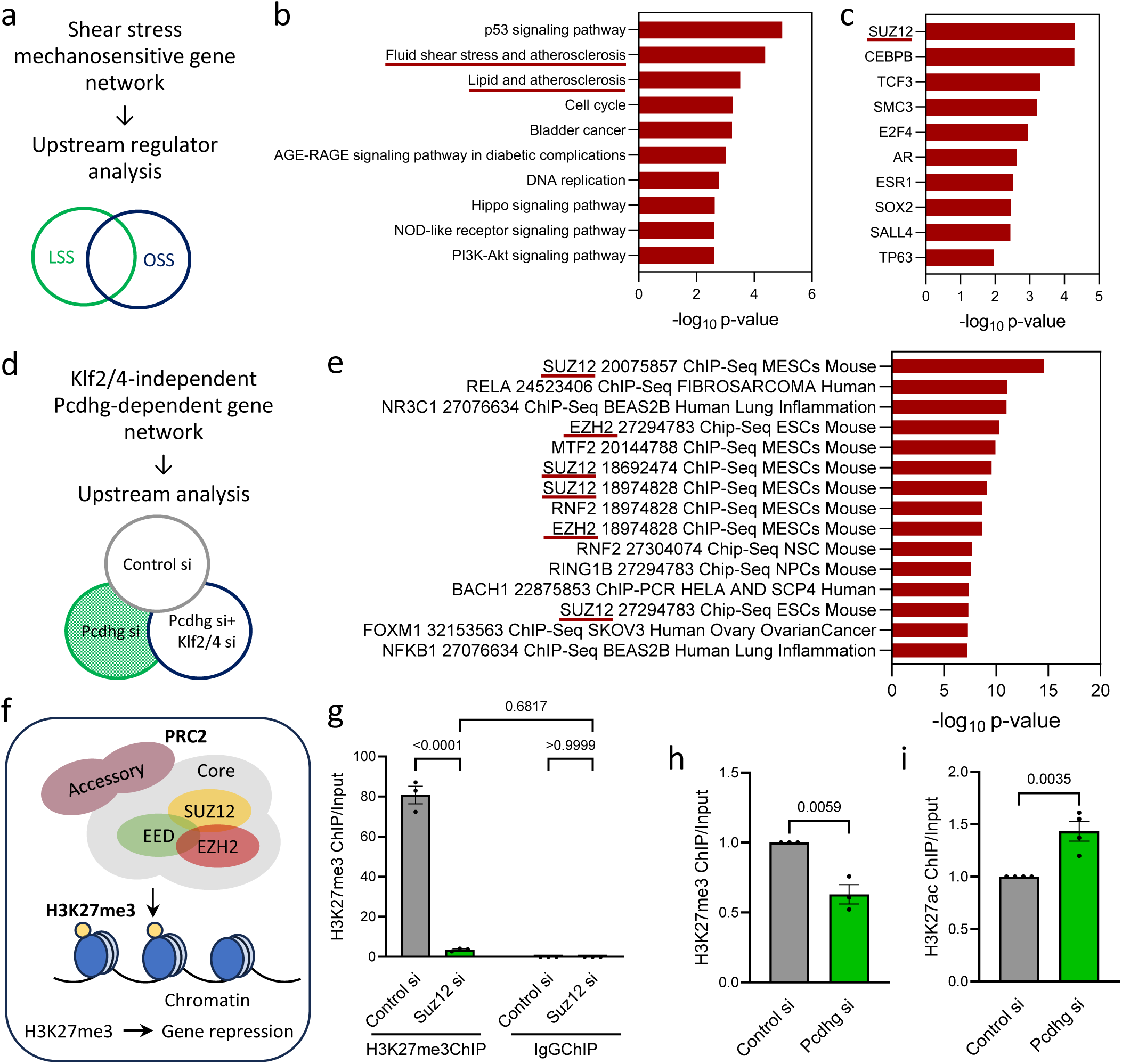
PRC2 in endothelial responses to flow. (a) Schematic of bulk RNAseq analysis of HUVECs exposed to LSS or OSS. (b, c) EnrichR analysis to identify enriched pathways/processes (b) and common upstream regulators of the DEGs (c) between LSS and OSS. (d) Schematic of bulk RNAseq analysis of Control si, Pcdhg si and Pcdhg+Klf2/4 triple si HUVECs to identify Pcdhg-dependent but Klf2/4 independent DEGs. (e) EnrichR analysis to identify common upstream regulators of the DEGs. (f) Schematic of PRC2 showing components and function. (g) H3K27me3 ChIP-qPCR at the Klf2 promoter. Depletion of PRC2 by Suz12 si was used as an internal control to confirm PRC2 specificity of H3K27me3 (N=3 independent experiments). (h, i) H3K27me3 and H3K27ac ChIP-qPCR analysis of Control si and Pcdhg si HUVECs on the Klf2 promoter (N=3 independent experiments). Values are means ± SEM. Statistical analysis used one-way ANOVA (h) or unpaired two-tailed Student’s t-test (i).

We recently reported that Pcdhg potently suppresses Klf2/4 in ECs^9^ (Fig 1d). We therefore assessed ECs after <u>Pcdhg si</u> as an independent approach. This experiment included <u>Pcdhg si + Klf2/4 si</u> conditions to distinguish direct Pcdhg targets from those affected through Klf2/4. EnrichR analysis of upstream regulators of the observed DEGs also identified EZH2 and SUZ12, two core components of PRC2, as the most significant hits (Fig 1e). Of the 494 DEGs (311 up and 183 down after Pcdhg suppression), 161 were identified as PRC2 targets, two-thirds (110/161) of which were upregulated after Pcdhg si while one-third (51/161) were downregulated. As PRC2 is mainly a transcriptional repressor, these data indicate that Pcdhg activates PRC2. This analysis also identified NF-κB, a known inhibitor of Klf2/4, and the PRC1 component RING1B, further validating the approach (Fig 1e). The PRC2 component Jarid2 was also identified as a suppressor of Klf2 in our original genome-wide CRISPR screen, further supporting this hypothesis^9^. Analysis of these DEGs using X2KWeb^11^, another tool for upstream regulator analysis, similarly showed PRC2 as a principal regulator of Pcdhg function (Fig S1b). However, transcripts of PRC2 components were unaffected by Pcdhg si, ruling out a feedback loop (Fig S1c). Interestingly, EZH2 protein was significantly reduced with a concomitant increase in Klf4 upon Pcdhg si, suggesting an effect on PRC2 complex stability^12^ (Fig S1d).

PRC2 is a protein complex comprising a core of EZH2, SUZ12, EED and RbAp46/48, plus accessory factors. EZH2 is the catalytic subunit of PRC2, which trimethylates lysine-27 on Histone H3 (H3K27me3) in promoters and enhancers of target genes to repress transcription (Fig 1f). ChIP-qPCR with an antibody to H3K27me3 pulled down the Klf2 promoter, whereas control IgG did not, but this signal was lost upon Suz12 si, validating Klf2/4 as direct targets of PRC2 in ECs (Fig 1g). Analysis of H3K27me3 ChIPseq datasets in HUVECs from the UCSC Genome Browser also showed peaks for H3K27me3 on the Klf2/4promoters (marked by nearby peaks for H3K27ac) (Fig S1e). As expected, ChIP-seq with an antibody to H3K27Ac, an activating mark that is mutually exclusive with H3K27me3, showed that Pcdhg si significantly increased H3K27ac at the Klf2 promoter (Fig S1f). ChIP-qPCR confirmed that Pcdhg si decreased H3K27me3 and increased H3K27ac on the Klf2 promoter (Fig 1h, i). Pcdhg suppression of Klf2/4 thus correlates with PRC2 epigenetic modification of their promoters.

### Elevated PRC2 activity in ASCVD suppresses protective Klf2/4

We next examined mouse and human arteries to address epigenetic H3K27 modification in vivo. In mice, the aortic arch contains adjacent regions under protective LSS (greater curvature) and disease-prone DSS (lesser curvature) where inflammatory molecules such as Vcam1, Icam1 are expressed in the endothelium^13^. *En face* staining of this tissue revealed high H3K27me3 in the lesser curvature relative to the greater curvature (Fig 2a), correlating PRC2 activity with disease susceptibility.

**Figure 2.**
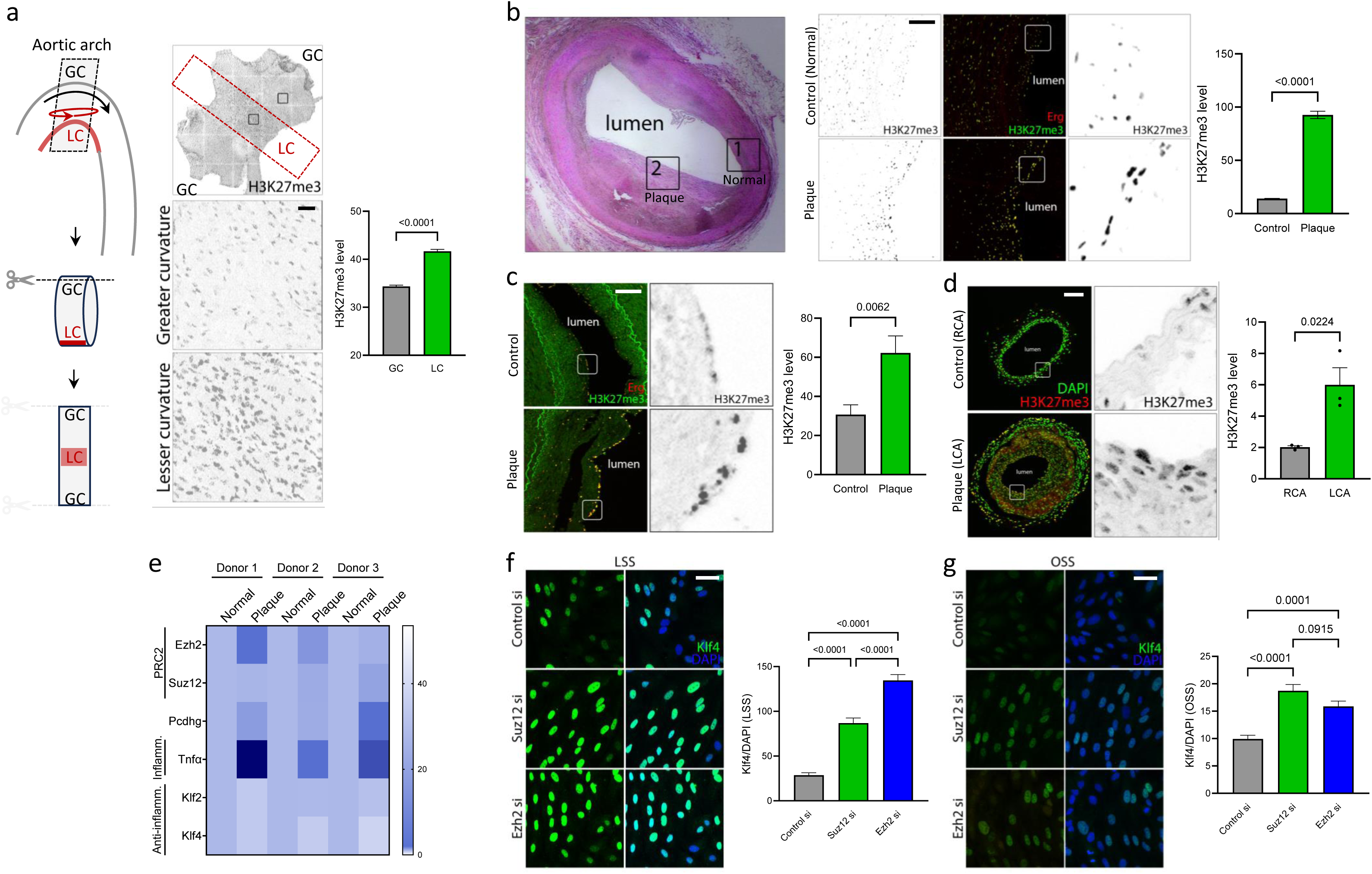
Elevated PRC2 activity in ASCVD suppresses Klf2/4. (a) Aortic arch region prepared en face and immunostained for H3K27me3 as a marker for PRC2 activity. Greater curvature (GC) is under LSS, lesser curvature (LC) is under DSS as shown in the schematic (N=3 animals). Graph: quantitation of H3K27me3 MFI. (b) De-identified human coronary artery sections from elderly donors stained with H&E (left) and H3K27me3 levels (right), comparing regions of plaque (Plaque region) to the regions of the same artery without evident plaque (Normal region). Sections were counter stained with Erg to mark ECs (N=3 donors). Graph: quantitation of HK27me3 signal intensity in ECs. (c) Artery sections from deidentified human CVD patients containing atherosclerotic plaques and non-diseased controls stained with H3K27me3. Sections were counter stained with Erg to mark ECs (N=8 donors). Graph: quantitation of H3K27me3 staining intensity in ECs. (d) Carotids from mouse Partial Carotid Artery (PCA) Ligation model of accelerated atherosclerosis stained for H3K27me3. Sections were counter stained with DAPI to mark nuclei (N=3 animals). RCA: Right Carotid Artery (control), LCA: Left Carotid Artery (plaque). Graph: quantitation of H3K27me3 staining intensity. (e) Re-analysis of Ezh2, Suz12 and Pcdhg in ECs from published scRNAseq data from deidentified human ASCVD patients comparing regions of plaque (Plaque) to an adjacent plaque-free region of the same artery (Normal)^14^. TNFα and Klf2/4 were used as internal controls for plaque ECs and normal ECs respectively (N=3 donors). (f, g) Depletion of PRC2 by Suz12 si or Ezh2 si in HUVECs exposed to LSS (f) or OSS (g) and stained for Klf4 (N=3 independent experiments). Graph: quantitation of Klf4 staining intensity. Values are means ± SEM. Statistical analysis used unpaired two-tailed Student’s t-test (b, c, d) or one-way ANOVA (f, g). Scale bar: (a) 50 μm, (b, c, d) 100 μm, (f, g) 20 μm.

Staining human arteries from deidentified elderly donors without symptomatic disease for H3K27me3 showed 4-fold higher H3K27me3 staining in plaque ECs compared to nearby unaffected regions of the same arteries (Fig 2b). Arteries from deidentified human ASCVD patients also showed a ∼3-fold increase in intimal H3K27me3 compared to control donors (Fig 2c). Plaques from a mouse model of ASCVD (Partial Carotid Artery Ligation and HFD for 3 weeks) again showed ∼3-fold higher staining for H3k27me3 (Fig 2d). These stains localized to both luminal ECs and to cells deeper in the vessel wall that were positive for lymphocyte common antigen, CD45 (Fig S2a). H3K27me3 is thus elevated in multiple instances of ASCVD, in ECs as well as leukocytes in the vessel wall, concomitant with increased expression of PRC2 components in ECs and immune cells.

Examination of published scRNAseq datasets from human atherosclerotic plaques also showed increased PRC2 and Pcdhg expression in plaque endothelium, compared to the control adjacent region of the same artery^14^ (Fig 2e). TNFα levels correlated positively, while Klf2 and Klf4 levels correlated negatively with ASCVD, serving as internal inflammatory and anti-inflammatory controls, respectively. This trend was less obvious in immune cells and SMCs (Fig S2b). Thus, tissue staining, published scRNAseq data and GWAS studies confirm upregulation/activation of PRC2 in plaque endothelium and perhaps other cell types.

To test whether PRC2 regulates Klf2/4 under flow, ECs transfected with control, Suz12, or Ezh2 siRNA were examined under LSS and OSS. We assessed Klf4 by immunostaining due to the availability of high-quality antibodies that are lacking for Klf2. Suz12 si and Ezh2 si elevated Klf4 in ECs under both flow patterns (Fig 2f, g). PRC2 thus suppresses Klf2/4 in ECs.

### Pharmacological inhibition of PRC2

PRC2 comprises methyltransferases EZH1/EZH2 along with other core and accessory components. EZH2 has much higher methyltransferase activity, is expressed at higher levels (Fig S3a, b) and its depletion strongly reduces H3K27me3 (Fig S3c), confirming it as the major H3K27 methyltransferase in ECs. A specific EZH2 inhibitor, Tazemetostat or EPZ-6438 (35-fold stronger inhibition of EZH2 than EZH1, and ∼100 fold stronger than other methyltransferases) is an FDA-approved drug used in cancer therapy^15^. Treatment with Tazemetostat (Tz) increased Klf4 in HUVECs under LSS (Fig 3a) compared to the untreated controls. We next examined the effects of Tz on EC inflammatory activation in vitro. HUVECs were treated with or without Il1β and Tz. Induction of VCAM1, a leukocyte adhesion receptor implicated in atherosclerosis, was strongly suppressed by Tz (Fig 3b). VCAM1-dependent adhesion of THP-1 monocytes to ECs that were activated by OSS was also potently blocked by Tz (Fig 3c). EZH2 methyltransferase activity thus suppresses Klf2/4 to permit vascular inflammation.

**Figure 3.**
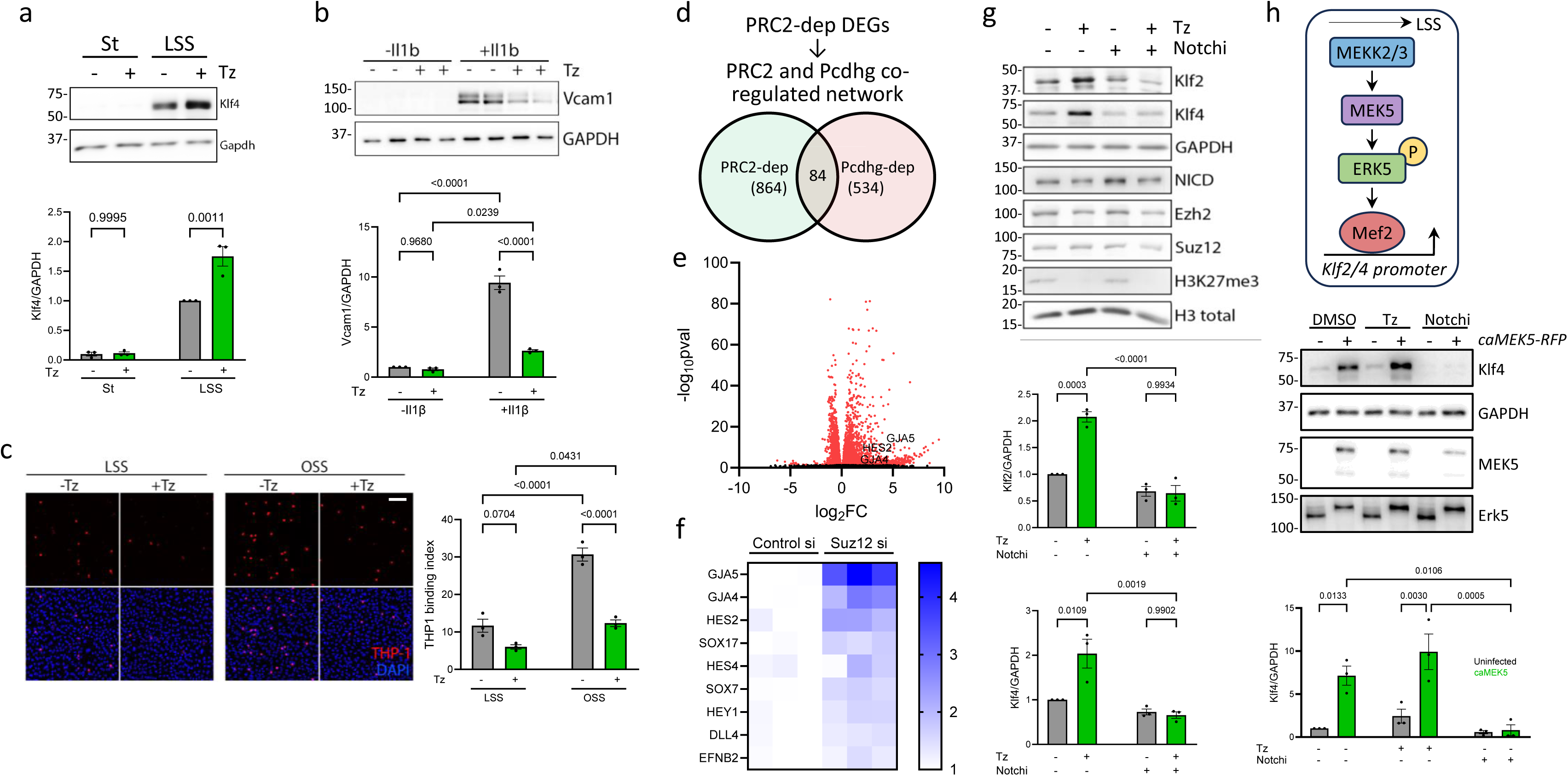
PRC2-Notch-Mek5 axis regulates Klf2/4. (a, b) Pharmacological inhibition of EZH2 catalytic activity with Tz in HUVECs under static (St) or LSS, immunoblotted for Klf4 and GAPDH loading control (N=3 independent experiments). Graph: quantitation of Klf4 normalized to GAPDH. (b) Immunoblot for VCAM1 in HUVECs, untreated or treated with Il1β (1 ng/ml) for 16 h with or without Tz (N=3 independent experiments). Graph: quantitation of VCAM1 normalized to GAPDH. (c) THP-1 monocyte binding to HUVECs under St or OSS for 16 h with or without Tz (N=3 independent experiments). Graph: quantitation of THP1 monocytes per field per condition (THP1 binding index). (d) Common processes/pathways regulated by PRC2 and Pcdhg. (e) Volcano plot from bulk RNAseq analysis of Control si and Suz12 si HUVECs to identify PRC2-regulated genes. (f) Heat map of Notch pathway targets in ECs upon Suz12 si compared to Control si from the RNAseq analysis. (g) Notch pathway was blocked by pharmacological inhibition of Notch-RBPJ interaction with RIN1 (Notchi) in HUVECs under LSS for 16h, with or without EZH2 inhibition by Tz, and immunoblotted for proteins, as indicated (N=3 independent experiments). Graphs: quantitation of Klf2 and Klf4 normalized to GAPDH loading control. (h) Schematic showing the Mekk2/3-Mek5-Erk5-Mef2 pathway. The pathway was activated by expression of caMek5, with or without PRC2 inhibitor Tz or Notch inhibitor RIN1, and immunoblotted for proteins, as indicated (N=3 independent experiments). Graph: quantitation of Klf4 normalized to GAPDH loading control. Values are means ± SEM. Statistical analysis used one-way ANOVA (a, b, c, g, h). Scale bar: (c) 100 μm.

### Increased Notch transcriptional activity upon PRC2i mediates Klf2/4 transcriptional activation

To understand how inhibiting PRC2 increases Klf2/4, we performed RNAseq analysis of control or Suz12 si HUVECs (Fig 3d-f). Of the DEGs after Suz12 si, 757 (87.6%) were upregulated while 107 (12.4%) were downregulated, consistent with transcriptional silencing by PRC2. Since PRC2 is downstream of Pcdhg, we searched for common transcriptional processes regulated by both PRC2 and Pcdhg (Fig 3d). Notch pathway targets were strongly upregulated in both Suz12 si and Pcdhg si (Fig 3e, f), which fits well with our previous finding that Notch is critical for Klf2/4 transcription^9^. Notably, Klf2/4 promoters contain both Notch-RBPJ binding sites^9^ and PRC2-deposited H3K27me3 (Fig S1e), suggesting opposing co-regulation by PRC2 and Notch.

We next assessed whether upregulation of Klf2/4 after inhibiting PRC2 required Notch transcriptional activity. Notch receptors mediate gene transcription after proteolytic cleavage and release of the intracellular domain (NICD), which translocates to the nucleus and binds the transcription factor RBPJ to activate gene transcription^16^. RIN1 (Notchi), a small molecule inhibitor of the NICD-RBPJ interaction that blocks Notch-dependent transcription, prevented Klf2/4 upregulation by Tz, without affecting the levels of PRC2 components (Fig 3g). Thus, PRC2 inhibition increases Notch-dependent transcription of Klf2/4.

### Notch functions via the Mekk2/3-Mek5-Erk5 pathway to induce Klf2/4

The best characterized mediator of Klf2/4 induction by LSS is the Mekk2/3-Mek5-Erk5 kinase cascade, which activates Mef2 transcription factors to directly induce Klf2/4 (Fig 3h)^17^. We also found that Notch directly induces Klf2/4 transcription^9^. Hence, to understand the role of PRC2, we designed an experiment to address the relationship between these players. We specifically activated the Erk5-Mef2 pathway by expressing doxycycline-inducible constitutively active Mek5 (caMek5)^18^ in HUVECs with or without inhibition of Notch (Notchi) and PRC2 (Tz). caMEk5 expression increased Klf4 ∼8-fold which was enhanced ∼2x by Tz and completely abolished upon Notchi (Fig 3h). Controls showed equivalent Mek5 levels and Erk5 activation under relevant conditions, and reduction of H3K27me3 by Tz. Klf4 is thus regulated via an integrated network in which PRC2 antagonizes the Mekk2/3-Mek5-Erk5-Mef2 cascade, which also requires Notch.

### PRC2 inhibition by Tazemetostat inhibits ASCVD in mice

Regulation of Klf2/4 via a PRC2-Mef2-Notch network implies that PRC2 inhibition or Notch activation will boost Klf2/4 expression and protect from ASCVD. Notch however is a widely expressed, multifunctional regulator, thus, is an unlikely target. However, the specific PRC2 inhibitor, Tazemetostat (Tz), is FDA approved, orally bioavailable and well tolerated for long-term in humans with limited adverse side-effects (ClinicalTrials.gov ID NCT02875548). We therefore examined Tz in a mouse model of ASCVD using an interventional strategy to mimic human clinical practice^19,20^. *Apoe^-/-^*mice were maintained on a high fat diet (HFD) for 12 weeks to induce hypercholesterolemia and ASCVD, then switched to chow diet with or without Tz. Vehicle or Tz was then administered in food for 6 additional weeks (Fig 4a). Tz treatment slowed plaque growth by >50% (Fig 4b). However, detailed characterization of plaques at the aortic root and brachiocephalic artery showed ∼2x thicker fibrous caps despite being smaller and essentially no expansion of necrotic cores (Fig 4c, d), a >2x reduction in CD68+ monocyte/macrophages (Fig 4e, f) and similarly reduced MMP9 levels (Fig 4g). Body weight and circulating lipids (TAGs and Cholesterol) were similar between vehicle and Tz treated cohorts, with expected reduction in Cholesterol upon switching to chow diet (Fig S4a-c). Tz interventional treatment thus moderately reduces plaque size but dramatically reduced markers of plaque vulnerability^21^. These data confirm PRC2 as a pro-inflammatory mediator in ASCVD and suggest possible use of Tz for treating advanced ASCVD.

**Figure 4.**
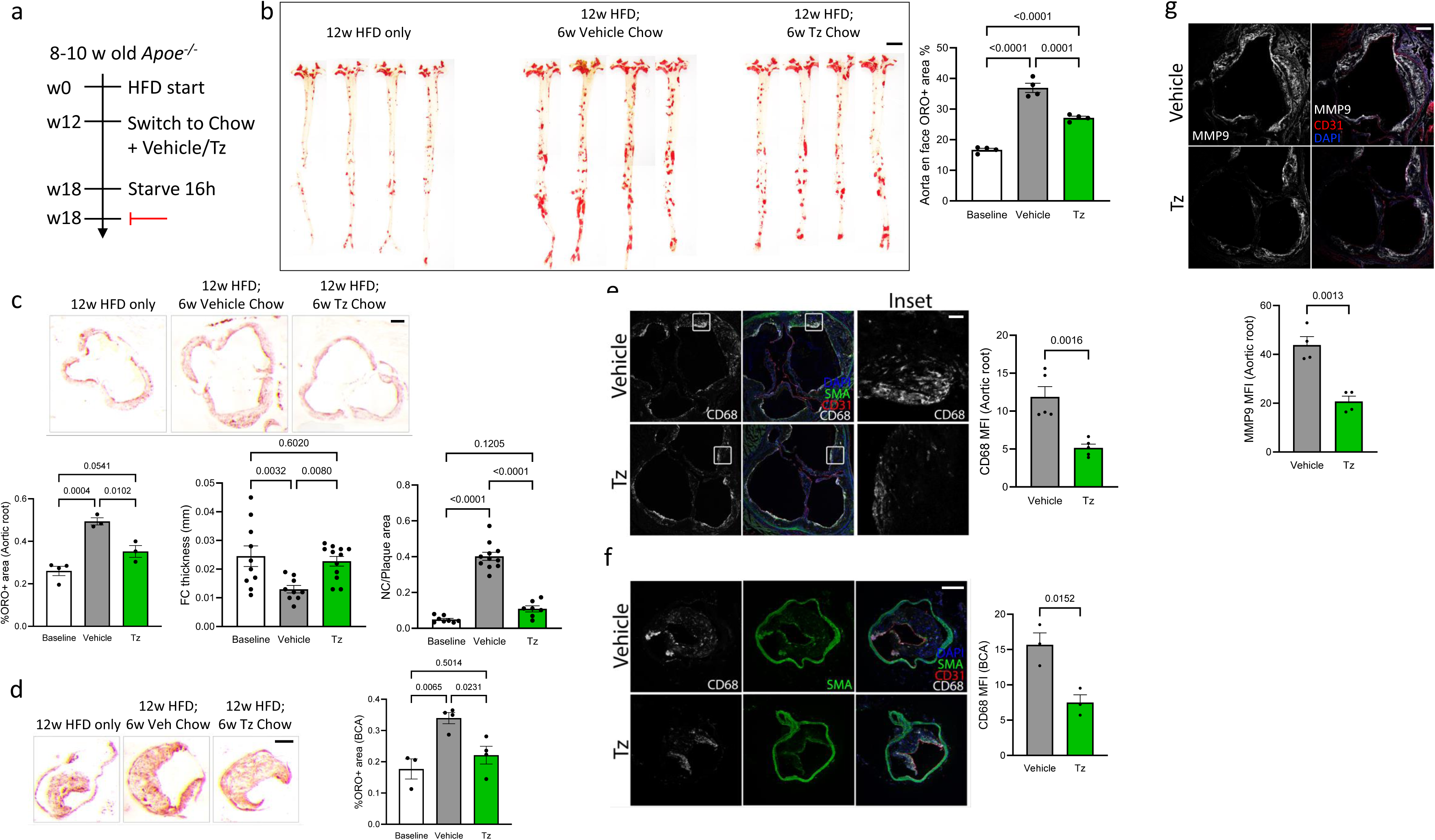
Interventional Tazemetostat treatment stabilizes plaque. (a) Schematic of experimental design. 8-10-week-old *Apoe^-/-^* mice were fed a High Fat Diet (HFD) for 12 weeks, switched to chow diet containing vehicle or Tz for an additional 6 weeks, and vessels analyzed. (b) Whole aortas were stained with Oil Red O to mark lipid-rich regions and imaged *en face* (N=4 animals). Graph: quantitation of percent Oil Red O positive area. (c, d) Sections from aortic roots (c) and brachiocephalic artery (BCA) (d) were stained with Oil Red O (N=3 animals). Graphs: quantitation of atherosclerotic plaque area (per animal), necrotic core (NC) area (per plaque) and fibrous cap (FC) thickness (per plaque). (e, f) Atherosclerotic plaques in aortic root sections (e) and BCA sections (f) were stained for the macrophage/monocyte marker CD68, and counterstained with SMA and DAPI to mark SMCs and nuclei respectively (N=3 animals). Graph: quantitation of CD68 MFI. (g) Aortic root sections were stained for plaque vulnerability marker MMP9, and counterstained with CD31 and DAPI to mark ECs and nuclei respectively (N=4 animals). Graph: quantitation of MMP9 MFI. Values are means ± SEM. Statistical analysis used one-way ANOVA (b, c, d) or unpaired two-tailed Student’s t-test (e, f, g). Scale bar: (b) 500 μm, (c, d) 100 μm, (e, f, g) 200 μm.

## 3. Discussion

Endothelial inflammatory activation underlies not only ASCVD but additional cardiovascular disorders including pulmonary arterial hypertension, acute lung injury and vascular dementia, thus, is of major relevance to human health^3,5,22–24^. Two independent bioinformatic analyses pointed toward PRC2 as a positive regulator of endothelial inflammatory activation. We found that PRC2 suppresses transcription of Klf2/4 by maintaining histone3 Lysine27 trimethylation in their promoters. Notch signaling, which is essential for expression of these genes, induces demethylation of Klf2/4 genes, allowing their transcription. In hyperlipidemic mice, the EZH2 inhibitor Tz slows progression of ASCVD and drastically improves plaque phenotype, providing in vivo support for this model.

PRC2 is best known for preserving the pluripotent state of stem cells and cancer cells by suppressing genes that promote differentiation^25^. Whether or how pro-inflammatory effects in ECs relate to this function is an open question. Studies of developmental and postnatal angiogenesis showed that arterial ECs are the last to form, originating from capillary and/or venous lineages, with no flow in the opposite direction^26–28^. We suggest that this arterial EC state is comparatively resistant to inflammatory activation and leukocyte recruitment. This notion is consistent with postcapillary venous endothelium as the main site of physiological leukocyte trafficking into tissues^29^. Both Notch activation and Klf2 expression are substantially higher in arterial ECs compared to capillary or venous. Re-analysis of published scRNAseq data from mouse arteries shows an inverse correlation between expression of inflammatory genes and arterial identity genes^14^ (Fig S2c, d). We therefore hypothesize that PRC2 functions in these two settings have common elements.

PRC2 is a general epigenetic suppressor of gene transcription that directly or indirectly targets thousands of genes^30^. Several studies showed the involvement of PRC2 in vascular inflammation and ASCVD^31^. The present study defines a mechanism of action and suggests possible clinical relevance. Importantly, most cardiovascular events are caused by plaque rupture, which correlates with the vulnerable plaque phenotype rather than plaque size^32–34^. We therefore utilized a “therapeutic protocol” in which *Apoe^-/-^* mice on HFD for 12 weeks with pre-existing disease were transferred to a chow diet to mimic lipid lowering therapy plus the EZH2 inhibitor Tz. Tz moderately slowed the rate of plaque growth (∼50%) but almost completely blocked the thinning of the fibrous cap, growth of the necrotic core and increase in immune cell content that identify plaque that is vulnerable to rupture. Tz is an FDA-approved drug used for long term treatment of hematopoietic cancers with activated EZH2 (ClinicalTrials.gov ID NCT02875548). In that setting, patients treated with Tz over an 8-year period showed no increase in deaths from other causes compared to placebo. Thus, testing in clinical trials for vascular inflammatory diseases including ASCVD may be feasible.

Major limitations of this study include general limitations of hyperlipidemic mice to model human disease, the likelihood of effects on gene targets in ECs other than Klf2/4 and possible effects of Tz on additional cell types. For this last point, PRC2 expression increases in plaque ECs is far more than in immune cells and negligibly in SMCs it may still play significant functional roles in these cell types. More detailed understanding of the molecular mechanisms governing Notch and PRC2-dependent transcription and examination of effects in vascular inflammatory diseases and are key areas for future work.

## 4. Materials and methods

### Primary cells, cell lines, and cell culture reagents

HUVECs pooled from 3 donors were obtained from the Yale Vascular Biology and Therapeutics Core. Cultures were screened for the absence of pathogens, maintained in EGM2 Endothelial Cell Growth Medium (Lonza, CC-3162) and used at passages 2-5. All cells were routinely screened for mycoplasma. SiRNA transfection was performed with Lipofectamine RNAiMAX (ThermoFisher, 13778150) in Opti-MEM medium (ThermoFisher, 31985070) using ON-TARGET plus SMARTpool siRNAs from Horizon Discovery (Dharmacon). For lentiviral transduction, HEK293T cells were transfected with lentiviral vectors along with pVSV-G (Addgene, #138479) and psPAX2 (Addgene, #12260) packaging plasmids using lipofectamine 2000 (ThermoFisher, 11668019) following the manufacturer’s instructions. Supernatants were collected 48-96 h after transfection and filtered through a 0.45 μm low-protein binding filter. Primary HUVECs were infected with lentivirus for 24h, then transferred to EGM2 medium. For monocyte adhesion assays, THP-1 cells labeled with CellTracker Deep Red (ThermoFisher, C34565) in HBSS supplemented with 1 mM Ca2+, 0.5 mM Mg2+, and 0.5% BSA were added to the HUVECs, incubated for 20 min at 37°C, washed 3x in HBSS and fixed with 3.7% formaldehyde. Cells were counterstained with DAPI before fluorescence imaging. Treatments used were Tazemetostat (Selleckchem), TNFα (PeproTech, 300-01A) and RIN1 (RBPJ Inhibitor-1) (Selleckchem, SS3376). All antibodies were validated in knockdown/knockout depletion by immunofluorescence and/or immunoblotting. Antibodies are listed in Table S1 and siRNAs in Table S2.

### Shear stress

Shear stress experiments were performed in parallel plate flow chambers and perfused within a pump and environmental control system as described^35^. Briefly, cells were seeded at 70-90% confluency on 10 μg/ml Fibronectin-coated glass slides for 48-72 h in complete medium. For shear stimulation, slides were mounted in custom-made 25×55 mm parallel plate shear chambers with 0.5 mm thick silicone gaskets and stimulated with 15 dynes/cm^2^ for LSS or 1 ± 4 dynes/cm^2^ for OSS in complete media. The media was maintained at 37°C and 5% CO_2_ with a heat gun and humidified bubbler, respectively.

### Animals and tissue preparation

All animal procedures followed Yale Environmental Health and Safety (EHS) regulations and were approved by Yale Institutional Animal Care and Use Committee (IACUC) and Yale Animal Resource Center (YARC). Mice were maintained in a light- and temperature-controlled environment with free access to food and water and all efforts were made to minimize animal suffering. Euthanasia was performed by overdose of inhaled isoflurane and death confirmed by subsequent cervical dislocation and/or removing vital organs and/or opening the chest cavity verified by the absence of cardiovascular function. Mice were perfused through the left ventricle with PBS and then 3.7% formaldehyde followed by tissue collection, as described^9^. The heart and spinal column, with aorta and carotids attached, were removed, fixed under gentle agitation for an additional 24 h at 4 °C, and washed 3x with PBS. For whole aorta *en face* prep, the isolated aortas were bisected along the lesser curvature and the aortic arch was bisected through the greater curvature as well. For aortic arch segment preparation, the aortic arch was bisected through the greater curvature. Hearts (containing aortic roots) and carotids were allowed to sink in 30% sucrose in PBS overnight at 4°C, embedded in OCT (optimal cutting temperature compound, Sakura, 4583) and frozen on dry ice before sectioning. Tissue blocks were cut into 8-10 μm sections using a cryostat (Leica) and sections stored at −80°C until use.

### Atherosclerosis progression and blood lipid analysis

To induce atherosclerosis, 8-12-week-old *Apoe^-/-^* mice were maintained on a high-fat diet (HFD; Clinton/Cybulsky high-fat rodent diet with regular casein and 1.25% added cholesterol; Research Diet, D12108c) for 12 weeks. Mice were then switched to chow diet containing either vehicle or Tazemetostat (Tz) at 150 mg/kg/day for a further 6 weeks. Tz amounts were calculated considering 3g food intake per mouse per day. Blood samples were collected in EDTA-coated tubes from overnight starved mice, centrifuged at 8,000g at 4°C for 10 min and the supernatant (plasma) was separated and stored at −80 °C. Total plasma cholesterol and triglycerides (TAG) were measured using kits according to the manufacturer’s instructions (Wako Pure Chemicals, Japan). Tissues were prepared and analyzed as described below.

### Tissue analysis

For IF, OCT tissue sections were thawed and washed 3x with PBS to remove OCT. Cells were fixed with 3.7% formaldehyde for 15 min at ambient temperature, washed with and stored in PBS. Deidentified human specimens were deparaffinized in Histo-Clear (National Diagnostics, HS-200). Sections were progressively rehydrated before antigen retrieval for 30 min at 95°C in 1X Antigen Retrieval Buffer (Dako, 51699). Samples were incubated in perm-block buffer (5% donkey serum, 0.2% BSA, 0.3% Triton X-100 in PBS) for 1 h at room temperature, incubated with primary antibodies in perm-block overnight at 4°C, washed three times in perm-block and then incubated with alexafluor-conjugated secondary antibodies (ThermoFisher) at 1:1000 dilution in perm-block for 1 h at room temperature. Slides were washed 3x in perm-block and 3x in PBS before mounting in DAPI Fluoromount-G (Southern Biotech; 0100–20). Images were acquired on a Leica SP8 confocal microscope equipped with the Leica Application Suite software. Confocal stacks were flattened by maximum intensity z-projection in ImageJ. Post background subtraction, the mean fluorescence intensity (MFI) or nuclear intensity (with DAPI mask) was recorded. For aorta ORO staining, the whole aorta was opened longitudinally on a soft-bottomed silica dish, incubated with 0.6% ORO in 60% isopropanol with gentle rocking for 1 h at ambient temperature, washed in 60% isopropanol for 20 minutes, washed in dH2O 3x, and mounted on slides with endothelium side up in OCT compound. Images were acquired with a digital microscopic camera (Leica DFC295). ORO staining on OCT tissue sections was done similarly. Quantitation of ORO positive area was done in ImageJ. Hematoxylin and eosin (H&E) staining on OCT tissue sections was done by Yale Research Histology Core using standard techniques. Plaque morphometric and vulnerability analysis was performed as described^9^. Plaque area was determined by ORO positive staining. For each plaque, the necrotic core (NC) area was defined as a clear area in the plaque that was H&E free, and the fibrous cap (FC) thickness was quantified by selecting the largest necrotic core and measuring the thinnest part of the cap.

### Immunoblotting

Cells were lysed in RIPA buffer (Roche) containing 1x Halt Protease Inhibitor Cocktail (ThermoFisher, 78429) and 1x PhosStop (Roche, 4906837001) for 30 min on ice, clarified at 13,000 g 4°C for 15 minutes, the supernatant transferred to new 1.5 ml tubes, 4x loading buffer (250mM Tris-HCl pH 6.8, 8% SDS, 40% glycerol, 20% β-mercaptoethanol, 0.008% bromophenol blue) added and the samples heated to 95°C for 5 min. Samples were resolved by a 4-15% gradient SDS Polyacrylamide gel, and transferred to 0.2 μm nitrocellulose membranes. Nitrocellulose was blocked with 5% non-fat skim milk for 1h at ambient temperature, incubated with desired antibodies diluted in 5% BSA using a standard immunoblotting procedure and detection by ECL (Millipore). Images were quantified in ImageJ by densitometry and normalized to GAPDH or Tubulin loading controls.

### RNA isolation, sequencing, and Chromatin Immuno-precipitation (ChIP), ChIP quantitative real-time PCR (ChIP-qPCR)

Total RNA was extracted from cells using the RNeasy Plus Mini Kit (Qiagen, 74136) according to the manufacturer’s instructions. RNA was quantified by NanoDrop, and RNA integrity was measured with an Agilent Bioanalyzer. Samples were sequenced using Illumina NovaSeq 6000 (HiSeq paired-end, 100 bp). The base calling data from the sequencer were transferred into FASTQ files using bcl2fastq2 conversion software (version 2.20, Illumina). PartekFlow (a start-to-finish software analysis solution for next generation sequencing data applications) was used to determine differentially expressed genes (DEGs). ChIP-qPCR primers are listed in Table S3.

### Quantification and Statistical Analysis

Morphometric analysis was done with ImageJ software (version 1.51; National Institutes of Health, Bethesda, MD). Graph preparation and statistical analysis was performed using GraphPad Prism 10.0 software (GraphPad software Inc.). Data were considered normally distributed, statistical significance was performed using Student t test for two group comparison or one-way ANOVA with Tukey’s post hoc analysis for multiple groups comparison, as described in the figure legends. Data are presented as means ± SEM. A p value less than 0.05 was considered significant.

## Data and Resource Availability

Data generated and/or analyzed during this study are freely available. Resources generated during this study are either freely available or available from the corresponding author upon reasonable request.

## Author contributions

DJ performed most of the experiments, analyzed data and prepared figures. DJ and RC performed and analyzed ChIP, ChIPseq, PCA ligation. DJ, TB performed Notch-Mek5 pathway analysis experiments. JF performed scRNAseq analysis. DJ, HD performed ASCVD experiments in mice. JGT Jr. and AWO provided human ASCVD samples. KAM supervised ChIP and ChIPseq experiments. DJ and MAS conceived the project, acquired funding for the project and wrote the manuscript. MAS supervised the project.

## Acknowledgments

The authors thank George Tellides for deidentified human elderly coronary artery samples, Sarah De Val for identification of Klf2/4 promoter sites, Dejian Zhao (Yale Center for Genome Analysis) for analysis of RNAseq data, Schwartz lab members for extensive discussions, Yale Center for Genome Analysis for RNAseq, and Yale Keck Oligo Synthesis Resource and DNA Sequencing Core. pSLIK MEK5-DD-mRFP1 neo was a gift from Kevin Janes (Addgene plasmid #47548; http://n2t.net/addgene:47548; RRID:Addgene_47548)^18^. This work was supported by National Institutes of Health grant R01 HL75092, a Leducq Trans-Atlantic Network Grant, and a Health Research consortium grant from Fundación Obra Social La Caixa (AtheroConvergence, HR20-00075) to MAS and American Heart Association grant #23POST1026109 to DJ.

---

**Figure S1.**
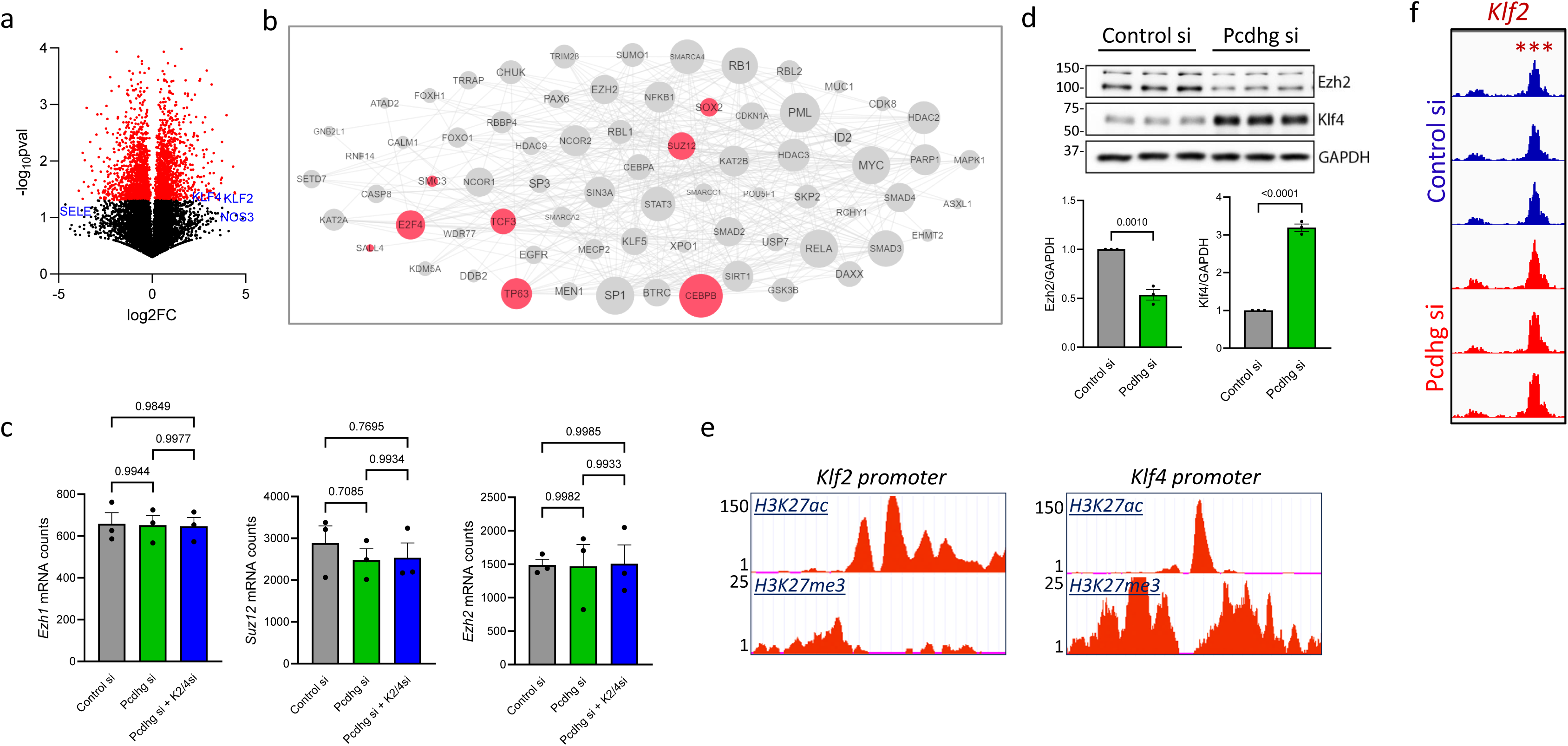
RNAseq and ChIPseq analysis. (a) Volcano plot from bulk RNAseq of HUVECs exposed to LSS vs OSS showing shear-regulated genes. (b) X2KWeb analysis to identify common upstream regulators of DEGs from bulk RNAseq analysis of Control si, Pcdhg si and Pcdhg+Klf2/4 triple si HUVECs. Red nodes represent transcription factors while gray nodes represent co-expressed DEGs. The size of the node represents centrality (number of connections with other nodes) in the network. (c) mRNA levels of Ezh2 and Suz12 after Pcdhg depletion. (d) Immunoblot for Ezh2 and Klf4 in Control or Pcdhg si HUVECs under laminar shear stress (LSS) (N=3 independent experiments). Graph: quantitation of Ezh2 and Klf4 normalized to GAPDH loading control. (e) Klf2 promoter region showing peaks for H3K27ac (marking promoter) and H3K27me3 obtained from available ChIPseq analysis on HUVECs (UCSC Genome Browser). (f) H3K27me3 ChIPseq analysis on Control si and Pcdhg si HUVECs showing a significantly higher peak call on Klf2 promoter (N=3 independent experiments). Values are means ± SEM. Statistical analysis used unpaired two-tailed Student’s t-test (d).

**Figure S2.**
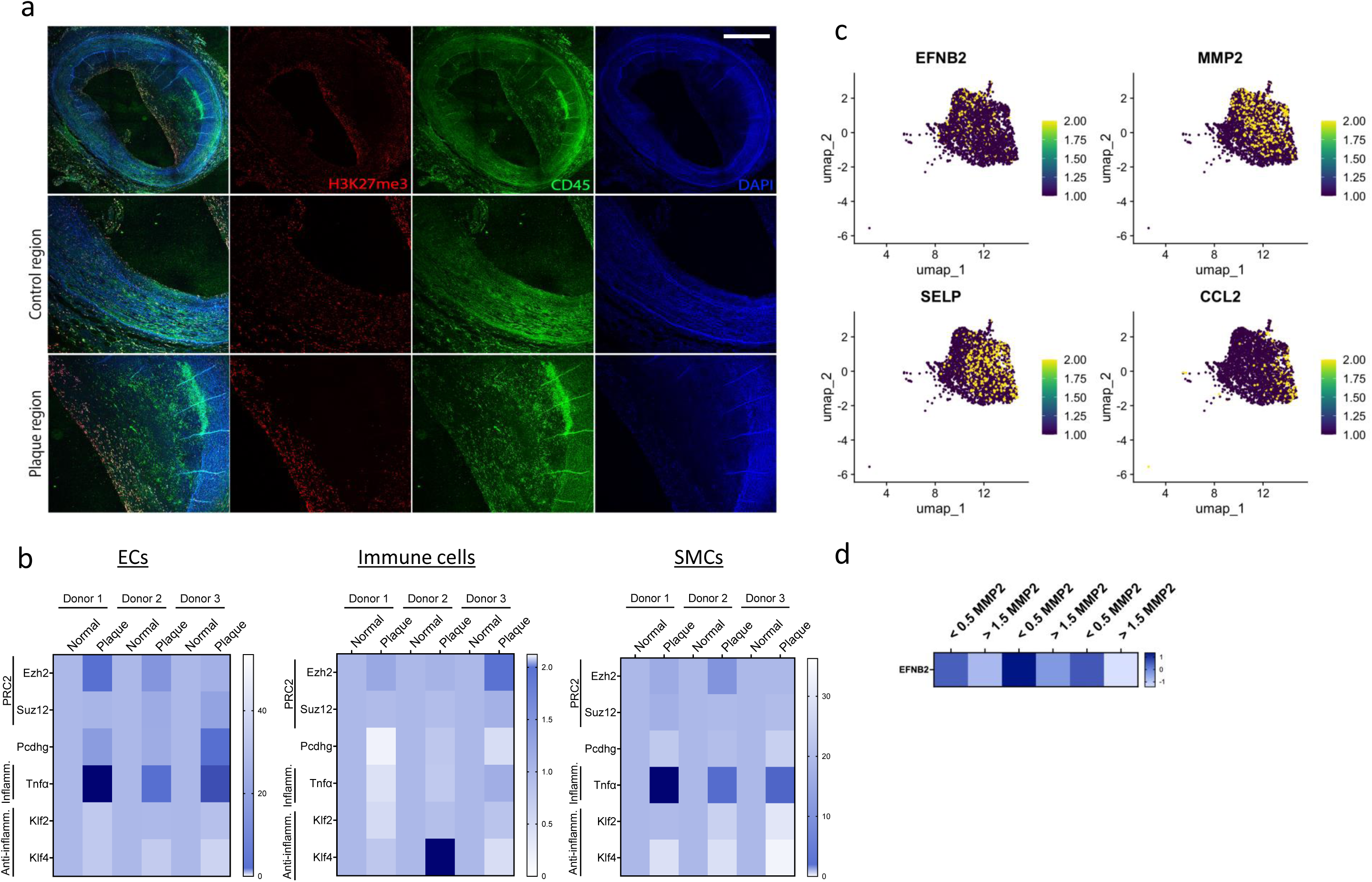
PRC2 is associated with CVD. (a) De-identified human coronary artery sections from elderly donors stained with H3K27me3 and leukocyte common antigen CD45, comparing regions of plaque (Plaque region) to the regions of the same artery without evident plaque (Control region). Sections were counter stained with DAPI to mark nuclei (N=3 donors). (b-d) Re-analysis of Ezh2, Suz12 and Pcdhg in major plaque cell types including ECs, immune cells and SMCs (b) from published scRNAseq data from deidentified human ASCVD patients comparing regions of plaque (Plaque) to an adjacent plaque-free region of the same artery (Normal)^14^. TNFα and Klf2/4 used as internal controls for plaque ECs and normal ECs respectively (N=3 donors). (c) Feature plot showing inverse correlation between the expression of arterial marker Efnb2 with inflammatory markers Mmp2, Selp and Ccl2 in plaque ECs. (d) Heat map showing Efnb2 levels in Mmp2^low^ (<0.5) vs Mmp2^high^ (>1.5) ECs from plaque.

**Figure S3.**
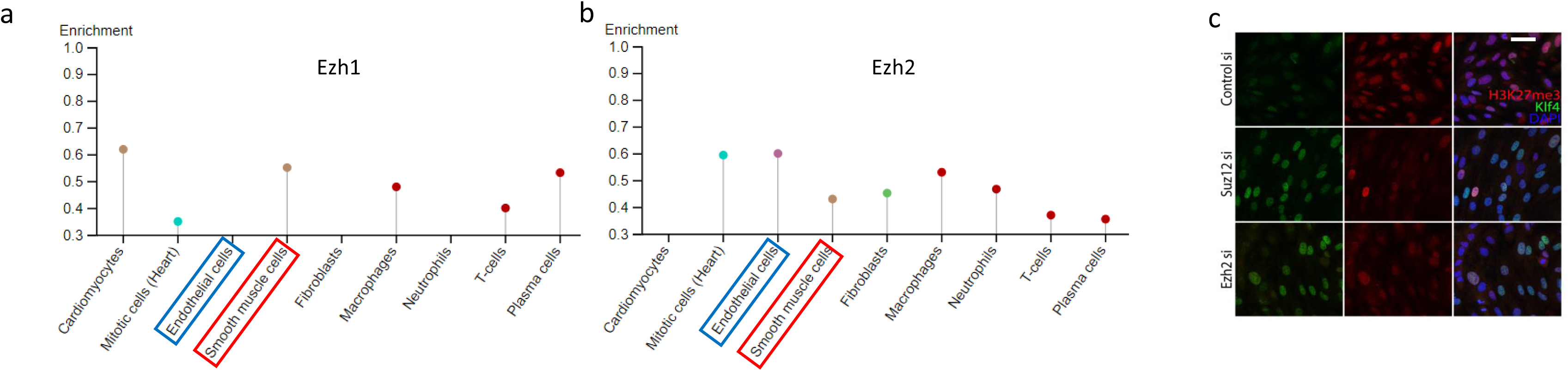
Expression of Ezh1 and Ezh2 in vascular cells. (a, b) Analysis of mRNA levels of Ezh1 and Ezh2 from human RNA atlas showing a comparison between ECs vs SMCs. Other relevant cell types shown, as indicated. (c) Functional validation of Ezh2 si and Suz12 si confirmed by depletion of PRC2 by Suz12 si or Ezh2 si in HUVECs exposed to OSS and stained for H3K27me3. Scale bar: (c) 20 μm.

**Figure S4.**
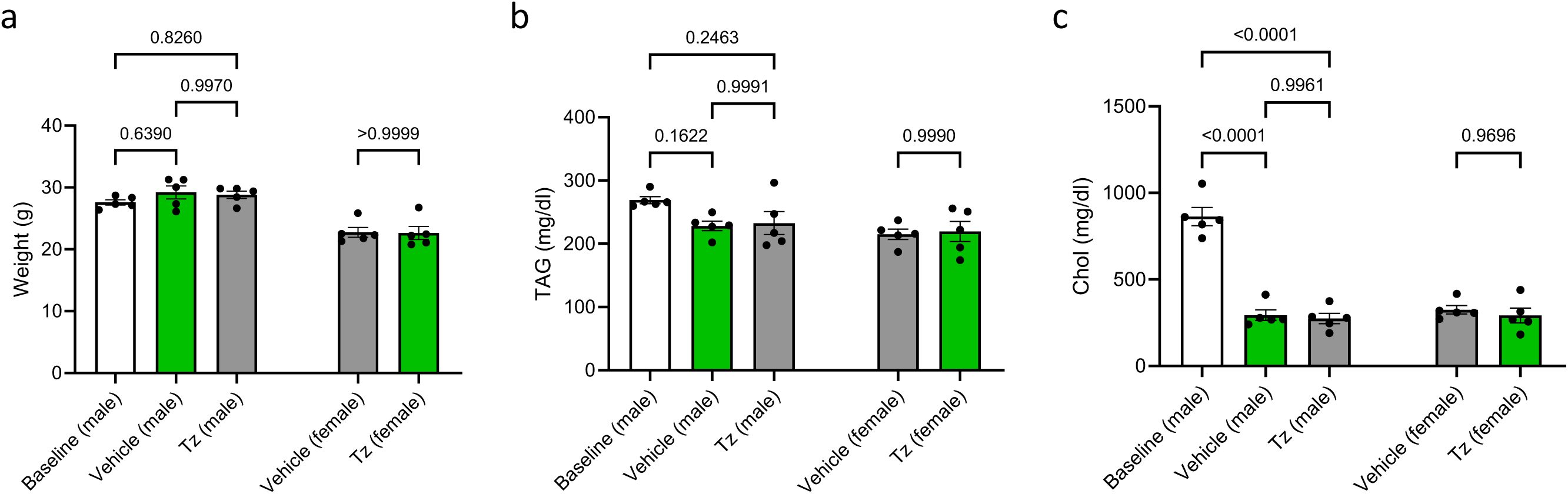
Mouse weight and blood lipid analysis. (a-c) Body weights (a), plasma triglycerides (TAGs) and cholesterol content of overnight starved mice treated as indicated (N = 5). Values are means ± SEM. Statistical analysis used one-way ANOVA.

**Supplementary Data Table S1.** List of primary antibodies.

| Antibody | Company, Catalog No. | Dilution for IF | Dilution for immunoblot |
| --- | --- | --- | --- |
| H3K27me3 | Cell Signaling, 9733 | 1:800 | 1:2000 |
| H3K27me3 | Abcam, ab6002 | 1:500 | 1:2000 |
| Klf4 | Abcam, ab215036 | 1:500 | 1:4000 |
| H3 total | Cell Signaling, 4499 | - | 1:2000 |
| Vcam1 | Abcam, ab134047 | - | 1:4000 |
| Klf2 | ProSci, 16-624 | - | 1:2000 |
| GAPDH | Cell Signaling, 5174 | - | 1:8000 |
| Ezh2 | Cell Signaling, 5246 | - | 1:2000 |
| Suz12 | Cell Signaling, 3737 | - | 1:2000 |
| NICD (Val 1744) | Cell Signaling, 4147 | - | 1:2000 |
| MEK5 | BD, 610956 | - | 1:2000 |
| ERK5 | Sigma Aldrich, E1523 | - | 1:4000 |
| MMP9 | Abcam, ab58803 | 1:400 | - |
| CD68 | ThermoFisher Scientific, 14-0681-82 | 1:200 | - |
| CD31 | R&D Systems, AF3628 | 1:200 | - |
| SMA | Millipore Sigma, A2547 | 1:500 | - |
| Erg | Abcam, ab92513 | 1:800 | - |
| CD45 | R&D Systems, AF114 | 1:200 | - |

**Supplementary Data Table S2.**
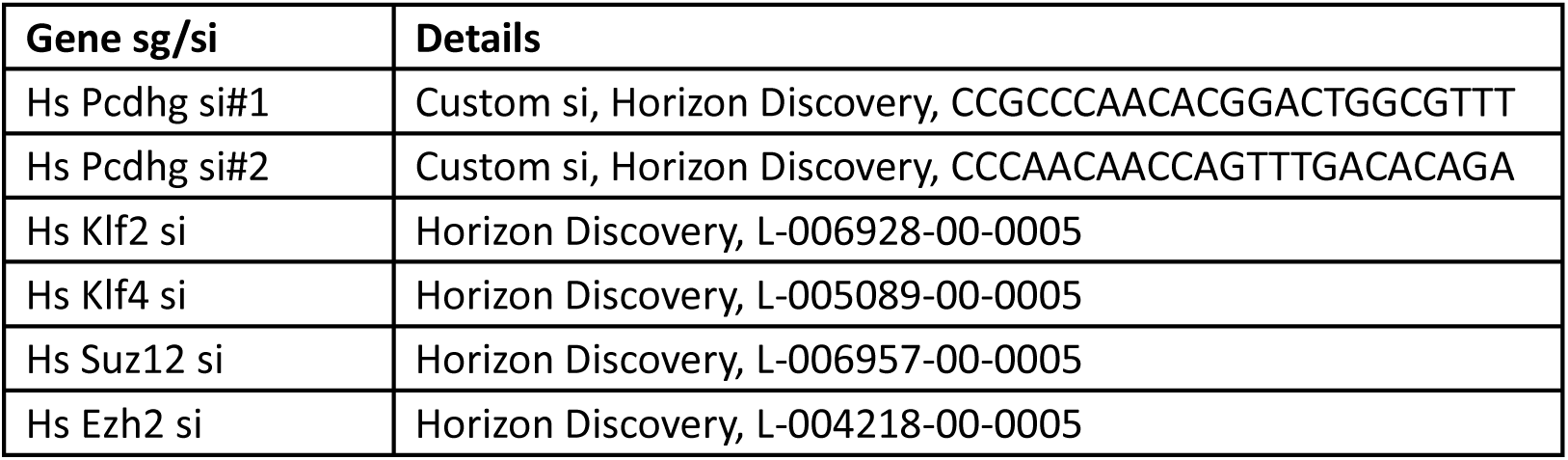
List of siRNAs.

**Supplementary Data Table S3.**
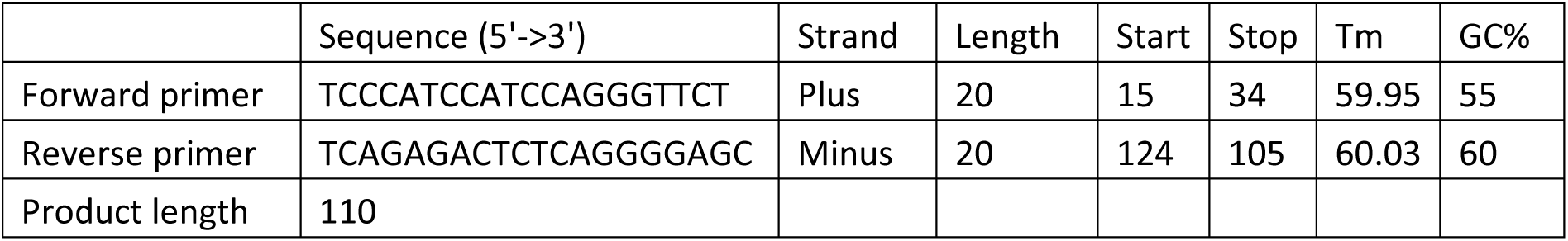
Hs *Klf2* promoter ChIP-qPCR primers.

